# Biphasic Npas4 expression promotes inhibitory plasticity and suppression of fear memory consolidation in mice

**DOI:** 10.1101/2023.11.17.567527

**Authors:** David V.C. Brito, Janina Kupke, Rostilav Sokolov, Sidney Cambridge, Martin Both, C. Peter Bengtson, Andrei Rozov, Ana M.M. Oliveira

## Abstract

Long-term memories are believed to be encoded by unique transcriptional signatures in the brain. The expression of immediate early genes (IEG) promotes structural and molecular changes required for memory consolidation. Recent evidence has shown that the brain is equipped with mechanisms that not only promote, but actively constrict memory formation. However, it remains unknown whether IEG expression may play a role in memory suppression. Here we uncovered a novel function of the IEG neuronal PAS domain protein 4 (Npas4), as an inducible memory suppressor gene of highly salient aversive experiences. Using a contextual fear conditioning paradigm, we found that low stimulus salience leads to monophasic Npas4 expression, while highly salient learning induces a biphasic expression of Npas4 in the hippocampus. The later phase requires NMDA receptor activity and is independent of dopaminergic neurotransmission. Our *in vivo* pharmacological and genetic manipulation experiments suggested that the later phase of Npas4 expression restricts the consolidation of a fear memory and promote behavioral flexibility, by facilitating fear extinction and the contextual specificity of fear responses. Moreover, immunofluorescence and electrophysiological analysis revealed a concomitant increase in synaptic input from cholecystokinin (CCK)-expressing interneurons. Our results demonstrate how salient experiences evoke unique temporal patterns of IEG expression that fine-tune memory consolidation. Moreover, our study provides evidence for inducible gene expression associated with memory suppression as a possible mechanism to balance the consolidation of highly salient memories, and thereby to evade the formation of maladaptive behavior.

## Introduction

The ability to convert environmental stimuli into long-term memory (LTM) is what allows an organism to adapt to the environment. While retaining novel information is essential for an individual’s life, the brain is also equipped with mechanisms that limit memory formation ^1–8^. This balance is thought to promote behavioral flexibility by preventing the formation of strong maladaptive memories that could impair the ability to adapt to changing environments ^1, 6^. However, how this is regulated at a cellular level is poorly understood.

It is well established that stimulus-induced transcription is essential for the consolidation of salient experiences into LTM ^9^. These inducible genomic signatures consist of temporally-defined waves of transcription of a broad functional repertoire of molecules in memory-associated regions, such as the hippocampus ^9, 10^. The earliest of these waves include the expression of immediate early genes (IEGs)^10^ and recently it has been suggested that experiences of distinct salience and valence generate unique IEG expression patterns ^11–18^. It remains however unknown whether stimulus salience triggers transcriptional responses that activate mechanisms that limit information storage, in addition to processes that favor memory consolidation. This could represent a biological strategy to prevent abnormally salient stimuli to give rise to strong memories that could compromise adaptive behavior.

In this study, we investigated the expression pattern of IEGs triggered by low and high salience fear conditioning paradigms. We found that highly salient experiences induce two phases of expression of the neuronal PAS domain protein 4 (Npas4) in the CA1 region of the dorsal hippocampus. Using pharmacological approaches, we demonstrated that the second phase of Npas4 transcription is dependent on N-methyl-D-aspartate receptor (NMDAR) activation. Intriguingly, inhibition of NMDAR activity, resulted in an enhancement of highly salient contextual fear memory, impaired extinction, and reduced contextual specificity of fear memory. Together this suggests a possible link between the second phase of Npas4 expression, suppression of consolidation of strong fear memories and facilitation of behavioral flexibility. In a complementary experiment, we demonstrated that time-specific exogenous Npas4 expression was sufficient to impair fear memory consolidation and improve contextual specificity of fear responses. Furthermore, we found that strong fear memory that induces biphasic Npas4 expression, as well as exogenous Npas4 expression were associated with increased inhibitory input from cholecystokinin-expressing interneurons (CCK^+^ IN) onto CA1 pyramidal neurons. Altogether, these findings suggest that besides its established positive role in memory formation ^7, 19^, Npas4 is an inducible memory suppressor gene that constrains memory consolidation of highly salient fearful experiences and favors behavioral flexibility, possibly to some extent through the modulation of CA1 inhibitory connectivity. Furthermore, our study reveals that experience salience is encoded by unique transcriptional signatures that include not only memory consolidation activators but also suppressors.

## Materials and Methods

### Mice

3-months-old C57BL/6N male mice (Charles River, Sulzfeld, Germany) were used for this study. Mice were group-housed (2-3 mice per cage), unless severe fighting occurred, and were housed on a 12h light/dark cycle with *ad libitum* access to water and food, 22 ± 1°C, 55 ± 10% relative humidity. All behavioral experiments took place during the light phase. Animals were kept singly-housed after cannula implantation to avoid cannula removal. Sick and/or injured mice from cage-mate fighting were excluded from this study. Animals were randomly assigned to experimental groups and blinded analysis was performed. All procedures were carried out in accordance with German guidelines for the care and use of laboratory animals and with the European Community Council Directive 86/609/EEC.

### qRT-PCR

Mouse dorsal hippocampal tissue was rapidly dissected and placed in RNAlater (Sigma, Munich, Germany) for 4 days at 4°C. Hippocampal tissue from home cage and fear conditioned mice was collected at the same time to account for possible time of the day effects. The CA1 region was microdissected and RNA was isolated using the RNeasy Plus Mini Kit (Qiagen, Hilden, Germany) with additional on-column DNase I digestion, according to the manufacturer’s instructions. RNA was reverse transcribed with the High-Capacity complementary DNA reverse-transcription kit (Applied Biosystems, Foster City, CA, USA) to generate complementary DNA. Quantitative reverse-transcription PCR (q-RT-PCR) was performed on a Step One Plus Real Time PCR System (Applied Biosystems, Foster City, CA, USA) using TaqMan gene expression assays (Applied Biosystems, Foster City, CA, USA) for the *Arc* (Mm00479619_g1), *c-Fos* (Mm00487425_m1) and *Npas4* (Mm00463644_m1). Expression levels of target genes were normalized to the expression of the housekeeping gene *GusB* (Mm00446953_m1). Controls were used to exclude the possibility of DNA or RNA contaminations.

### Western Blotting

Hippocampal cultures infected on day *in vitro* (DIV) 4 were lysed on DIV 10 in boiling SDS sample buffer (160 mM Tris-HCl (pH 6.8), 4% SDS, 30% glycerol, 10 mM dithiothreitol, and 0.02% bromophenol blue). In the case of western blotting of tissue samples, the dorsal CA1 was quickly microdissected in ice-cold phosphate-buffered saline (PBS) and homogenized in RIPA buffer (150 mM NaCl, 1% Triton X-100, 0.5% sodium deoxycholate, 0.1% SDS, 50 mM Tris, pH 8.0) supplemented with 1% protease inhibitor cocktail (Sigma-Aldrich, Munich, Germany). Protein concentration was measured by Bradford assay and 20 µg of protein was loaded in a 10% polyacrylamide gel after being denatured at 95°C for 5 min. After SDS-PAGE, gels were blotted onto a nitrocellulose membrane (GE Healthcare, Buckinghamshire, UK) and later blocked in 5% milk in PBS-T and probed with the following antibodies: Arc (1:6000, Synaptic Systems, #156003), Fos (1:1000, Cell Signaling, #2250), Npas4 (1:1000, Activity Signaling), α-Tubulin (1:400 000, Sigma #t9026) and β-Actin (1:1000, Santa Cruz, #SC-47778). Antibodies were diluted in 5% milk in PBS-T. Next, the membranes were incubated with horseradish peroxidase-conjugated secondary antibodies and later analyzed using a ChemiDoc^TM^ Imaging System (Bio-Rad, California, USA).

### Immunofluorescence

Intracardiac perfusions of deeply anesthetized mice were performed with ice-cold PBS and 4% paraformaldehyde (Sigma-Aldrich, Munich, Germany). Brains were collected and further post fixed in 4% paraformaldehyde (Sigma-Aldrich, Munich, Germany) overnight at 4 °C. Next day, brains were transferred to a 30% sucrose solution in PBS and left for 3 days. Brain slices were cut at a thickness of 30 μm. For the analysis of inhibitory synapses, slices were photobleached prior to immunofluorescence by incubation with 5% hydrogen peroxide (Sigma-Aldrich, Munich, Germany) for 2 h under white light at room temperature. After blocking in 8% normal goat serum with 0.3% Triton X-100 in PBS for 1 h at room temperature, slices were incubated with primary antibodies raised against the HA-tag (1:1000, Covance, MMS-101R), VGAT (1:500, Synaptic systems #131 011), gephyrin (1:1000, Synaptic systems #147 318) or CB1R (1:500, Synaptic systems #258 008) diluted in PBS containing 2% normal goat serum and 0.3% Triton X-100 at 4 °C overnight. Next day, after washing 3x for 5 min with PBS, slices were incubated with secondary antibodies (1:500, goat anti-guinea pig Alexa488; goat anti-rabbit Alexa597; goat anti-mouse Alexa647 [Life Technologies, Eugene, OR, USA]) diluted in the same solution as primary antibody for 2 h in the dark at room temperature. Lastly, after washing 3x for 5 min with PBS, slices were incubated in Hoechst 33258 (2 μg/ml, Serva, Heidelberg, Germany) for 5 min and mounted on glass slides. Synaptic analysis was based on Hartzell et al 2019 ^20^ with few alterations. Briefly, a background subtraction was applied to each channel of confocal acquired images. Next, the same threshold was applied across images of each channel from all conditions using the FIJI software^21^. The integrated density (the product of the area and mean grey value) that resulted from the overlap of the three fluorescent signals (VGAT, Gephryrin and CB1R), was quantified within regions of interest (ROIs) for superficial and deep layers of dorsal middle CA1 regions (CA1b ^22^). Superficial and deep CA1 regions were classified based on proximity to the *stratum radiatum* (superficial) or *stratum oriens* (deep). The integrated density was normalized to cell number within each ROI as determined by Hoechst counterstaining.

### Behavioral testing

Mice were habituated to the experimenter and behavioral room by daily handling for 3 consecutive days, 2 minutes per mouse. Contextual fear conditioning was performed as previously described ^23, 24^ with the following modifications. Two strong foot shock protocols were used to induce a biphasic wave of Npas4 expression. In one protocol mice were allowed to explore the conditioning chamber for 2 min and 28 s and three-foot shocks (0.7mA, 2s) were administered with 2 min and 28 s as inter-shock interval. After the last shock, the animals remained for 60 s in the chamber before returning to the home cage. In the second protocol, after an initial chamber exploration of 3x 2 min and 28 s, the mice received one foot shock (0.7 mA, 2 s). After the shock the animals remained in the chamber for 60s before returning to their home cage. A weak foot shock protocol that does not induce late Npas4 expression consisted in 2min and 28s of exploration until a 0.2 mA footshock was administered for 2 s. After the shock the animals remained for 30s in the chamber before returning to their home cage. The testing session consisted in exposing the animals to the conditioning chamber for 5 min in the absence of foot shocks. For the altered context testing, the context was highly dissimilar to the conditioning chamber. Specifically, it was located in a different experimental room and the properties of the chamber – floor (white plastic instead of metal grid), scent (lemon detergent instead of ethanol), shape (triangle instead of square) and the light intensity - were changed. The fear memory extinction training consisted in exposing the animals to the conditioning chamber for 5 min in the absence of foot shocks for 7 consecutive days starting after the contextual fear conditioning.

### Cannula implantation and infusion

Mice were implanted with 26-gauge double guide cannula cut 1mm below the pedestal (C235G-3.0/Spc, Plastics One, Bilaney). The target injection site coordinates (CA1 region of the dorsal hippocampus) were the following: − 2 mm anteroposterior, ± 1.5 mm medio-lateral, − 1.2 mm dorsoventral. Cannulas were placed using HY-bond polycarboxylate cement (9917-1, Shofu) and 2 screws (00-96×1/16, Plastics One, Bilaney) and left to dry for 25min. After this period, a dummy cannula without projection (c235g-3 Plastics One, Bilaney) was placed inside the guide cannula to avoid clogging. The animals were allowed to recover from surgery for 7 days before experiments. At the time of drug delivery an internal infusion cannula (C235G-3, Plastics One, Bilaney) was tightly fitted into the guides and injections (0.5μl/side) of DL-2-amino-5-phosphonopentanoic acid (DL-APV) (5μg/μl, 22.8mM), SCH23390 (5μg/μl, 15.4mM) or saline were performed at 200nl/min speed with a microinjection pump. The infusion cannulas were left in place for 60 additional seconds to minimize backflow. Cannula placement was verified *postmortem* during tissue microdissection. Only data from animals with correct placement were analyzed.

### Recombinant Adeno Associated Virus (rAAV) production

Viral particles were produced and purified as described previously ^25^. Briefly, rAAVs were produced by co-transfection of human embryonic kidney (HEK) cell line 293 (Stratagene, California, USA) with the target AAV plasmid and helper plasmids (pFΔ6, pRV1and pH21) using standard calcium phosphate precipitation. 60 h after transfection, HEK 293 cells were harvested and lysed. Finally, the viral particles were purified using heparin affinity columns (HiTrap Heparin HP; GE Healthcare, Uppsala, Sweden) and concentrated using Amicon Ultra-4 centrifugal filter devices (Millipore, Bedford, MA). The dual-component TetON-based system contains the driver plasmid (rAAV-hSyn-rtTA-T2A-TetR-KO) that expresses under the control of a neuron-specific promoter (hSynapsin), the transactivator (rtTA), the tetracycline repressor (TetR) and the fluorescent protein Kusabira Orange (KO) that serves as an infection marker^26^. In the second construct a human influenza hemagglutinin (HA)-tagged eGFP (rAAV-TRE-eGFP-HA) or full length Npas4 (rAAV-TRE-HA-Npas4) expression cassette is under the control of the tetracycline responsive promoter (TRE). For the generation of these vectors, the pAAV-PTRE-tight-hM3Dg-mCherry ^27^ (a gift from William Wisden [Addgene plasmid # 66795; http://n2t.net/addgene:66795; RRID:Addgene_66795]) was used. Specifically, HA-GFP or HA-Npas4 coding sequences were inserted into the pAAV-PTRE-tight-hM3Dg-mCherry after excision of the hM3Dq-mCherry transgene. For each virus batch produced, the infection rate, toxicity, viral titer was evaluated before the onset of experiments. Viral titers obtained after production of all viruses were similar and were matched to obtain final working concentrations of 10^12^ viral particles/mL.

### Stereotaxic surgery

rAAVs were injected into the dorsal CA1 at the following coordinates relative to Bregma: − 2 mm anteroposterior, ± 1.5 mm medio-lateral, − 1.2 mm dorsoventral. A total volume of 0.5 μl (1:1 ratio of Driver and eGFP or Npas4) was injected per hemisphere at 100 nl/min. Before and after injections at each individual site, the needle was left in place for 5 min. At the time of behavioral experiments, the experimenter was blind to the identity of the virus injected into each mouse. Behavioral experiments started 3 weeks after rAAVs delivery. Intraperitoneal doxycycline hyclate (2.5 mg in 500 µL saline solution, 100 mg/kg, Sigma-Aldrich, Munich, Germany) injections were performed immediately or 12 hours after contextual fear conditioning training. After behavior, histological analysis was performed to confirm correct targeting and tissue and cellular integrity. Mice that showed absence or miss targeting of viral expression were excluded.

### Primary hippocampal cultures

Hippocampal cultures from newborn C57Bl/6N mice (Charles River, Sulzfeld, Germany) were prepared and maintained as previously described ^25^. Briefly, mice hippocampi were dissociated at P0 by papain digestion and plated onto tissue culture dishes coated with poly-D-lysine and laminin (Sigma-Aldrich, Munich, Germany). The primary cultures were maintained for 8 days in Neurobasal-A medium (GibcoTM) supplemented with 1% rat serum (Biowest), 0.5mM L-glutamine (Sigma-Aldrich, Munich, Germany) and B27 (Gibco™), followed by incubation in salt-glucose-glycine solution (10 mM HEPES, pH 7.4, 114 mM NaCl, 26.1 mM NaHCO3, 5.3 mM KCl, 1 mM MgCl2, 2 mM CaCl2, 30 mM glucose, 1 mM glycine, 0.5 mM sodium pyruvate, and 0.001% phenol red) and phosphate-free Eagle’s minimum essential medium (9:1 v/v), supplemented with insulin (7.5 μg/ml), transferrin (7.5 μg/ml), and sodium selenite (7.5 ng/ml) (ITS Liquid Media Supplement, Sigma-Aldrich, Munich, Germany) and penicillin-streptomycin. rAAV infection of cultures occurred on DIV 4. Infection rates were determined by analyzing the respective transgene expression which ranged from 80-90%. Experiments were performed on DIV 10-11. DNA co-transfection was performed after a culturing period of 8 DIV using Lipofectamine 2000 (Invitrogen, CA, USA) as described previously ^23, 28, 29^. Doxycycline hyclate (25μM, Sigma-Aldrich) was introduced in the medium at DIV 8. N numbers represent independent cell preparations.

### Luciferase Assays

Assays were performed as previously described with minor alterations ^28^. We used the following Firefly Luciferase expression vectors: pGL4.29[luc2P/CRE/Hygro] (Promega) that contains four CREB responsive element (CRE) cis-elements and a minimal promoter (4×CRE-pmin), a Npas4 reporter plasmid contained four Npas4 responsive elements (NRE) (TCGTG), a consensus binding motif for Npas4 and a minimal promoter (kindly provided by Dr.Yingxi Lin ^30^). Additionally, the plasmid pGL4.83h[RlucP/Puro] (Promega) that contains the human EF1a promoter in upstream to the Renilla luciferase (Rluc) was used for normalization. On DIV 8, mouse primary hippocampal cultures in 48-well plates were changed to transfection medium. Lipofectamine 2000 (Invitrogen, CA, USA) was used for transfection according to the manufacturer’s instructions. Neurons were co-transfected with the following constructs: driver plasmid, eGFP or full length Npas4 and 4×CRE-pmin or 4×NRE-pmin (1 µg/well), together with Rluc (75 ng/well). DNA (µg): Lipofectamine 2000 (µl) ratio of 1:2 in total of 25 µl/well. On DIV 10, Dual-Glo Luciferase Assay System (Promega) was used to measure Firefly luciferase (FFluc) and Renilla luciferase (Rluc) activity levels. Background signal measured from non-transfected cells was subtracted and FFluc levels were normalized to Rluc levels. Each condition was performed in triplicate.

### Electrophysiological Recordings

Transverse hippocampal 300 μm acute brain slices were prepared from mice anaesthetized with CO2. The slicing chamber contained an oxygenated ice-cold solution [modified from ^31^] composed of (in mM): K-Gluconate, 140; N-(2-hydroxyethyl) piperazine-N′-ethanesulfonic acid (HEPES), 10; Na-Gluconate, 15; ethylene glycol-bis (2-aminoethyl)-N, N, N′, N′-tetraacetic acid (EGTA), 0.2; and NaCl, 4 (pH 7.2). Slices were incubated for 30 min at 35°C before being stored at room temperature (22– 24°C) in artificial CSF (ACSF) containing (in mM): NaCl, 125; NaHCO3, 25; KCl, 2.5; NaH2PO4, 1.25; MgCl2, 1; CaCl2, 2; and glucose, 25; bubbled with 95% O2 and 5% CO2.

Whole cell patch clamp recordings were done at 33-34°C from submerged slices with constant perfusion of ACSF. CA1 pyramidal neurons were visualized with a differential interference contrast. The liquid junction potential was calculated to be 15 mV and subtracted from the voltage data.

Tetrodotoxin (TTX, 1 μM) was added to the perfusion bath to block Na^+^-dependent action potentials. Miniature inhibitory postsynaptic currents (mIPSCs) were then recorded at 0 mV, the reversal potential of ionotropic glutamate receptors, to mask the presence of excitatory postsynaptic currents. Control mIPSCs were recorded for at least 10 minutes, before adding the CB1 receptor antagonist AM0251 (1 μM) and recording for an additional 20 minutes. Access resistance and series resistance were monitored before and after each recording. Cells with an increase of series resistance >15% during the experiment were discarded. For analysis, we used traces obtained within the last 5 minutes of control and AM-treated recordings. Data were analyzed with the Mini Analysis Program (Synaptosoft).

Data, depending on normality of data point distribution, are presented as mean ± SD or medians, 25th, 75th percentiles and individual values obtained in control and in the presence of AM-251 (1 μM). On the box plots whiskers show min and max values. Statistical analysis was performed using the paired T-test or the Wilcoxon Signed Rank Test within the SigmaPlot software (Systat, USA).

### Statistical analysis

Each set of experiments contained mice injected with control or experimental viruses and were randomized per cage (i.e., each cage of four mice contained mice injected with control or experimental viruses). Mice that underwent pharmacological CA1 infusions were individually housed. After stereotaxic surgery and until the end of each experiment, the experimenters were blind to the experimental group of each mouse. For normally distributed data sets, two-tailed unpaired Student’s t tests were used to compare two groups. If a dataset was compared more than once a one- or two-way ANOVA was used followed by appropriate multiple comparison *post hoc* tests to control for multiple comparisons as specified. In the case of a non-Gaussian distribution, two-tailed Mann-Whitney tests were used to compare two distinct groups, or a Kruskal-Wallis test followed by Dunńs *post hoc* test to compare more than 2 groups. The sample size was determined based on similar experiments carried-out in the past and in the literature. Except for electrophysiological analysis, all plotted data represent mean ± SEM. Statistical analysis was performed using GraphPad Prism, version 9 or SigmaPlot (Systat, USA). Variance between groups was not formally tested. For behavioral experiments the investigators were blind to group allocation during data collection and analysis. All behavioral sessions were video recorded, and an experimenter blinded to the group identity performed manual scoring to determine freezing behavior. The exact sample size is depicted in the respective figure legend and by individual dots in figures.

## Results

### High salience contextual learning induces biphasic Npas4 expression in the hippocampus

To address the molecular mechanisms associated with experience salience, we established two contextual fear conditioning protocols of low (1×0.2 mA) or high (3×0.7 mA) stimulus salience that induced weak or strong fear memory, respectively (Figure 1 A-B). Next, we monitored the expression of IEGs in mice that underwent learning in both protocols compared to homecage (HC) control mice (Figure 1 C) during the critical phase of memory consolidation ^32, 33^, as it is known that variation in salience of a learning stimulus affects IEG expression ^13, 15, 34^. Of note, tissue from HC controls and fear conditioned mice was obtained at the same time of the day to account for possible time of the day effects. Particularly, we analyzed the mRNA levels of the activity-regulated cytoskeleton-associated protein (Arc), c-Fos and Npas4 in the CA1 region of the dorsal hippocampus. As expected, we detected increased levels of IEG expression compared to HC animals 15’ to 1h after learning in animals that underwent low (Figure 1 D-F) or high (Figure 1 H-J) salience fear conditioning. Expression of Arc and c-Fos levels generally returned to baseline levels 4h after learning in both conditions (Figure 1 D-E, H-I). In low salience fear conditioning, Npas4 mRNA and protein levels also returned to baseline (Figure 1 F-G). However, mice that underwent high salience fear conditioning showed a late Npas4 expression at 4h (Figure 1 J), which was also detected at the protein level at 6h (Figure 1 K). These results indicated that high salience fear conditioning triggers a biphasic Npas4 expression, which is not present after a low salience stimulus is applied.

**Figure 1.**
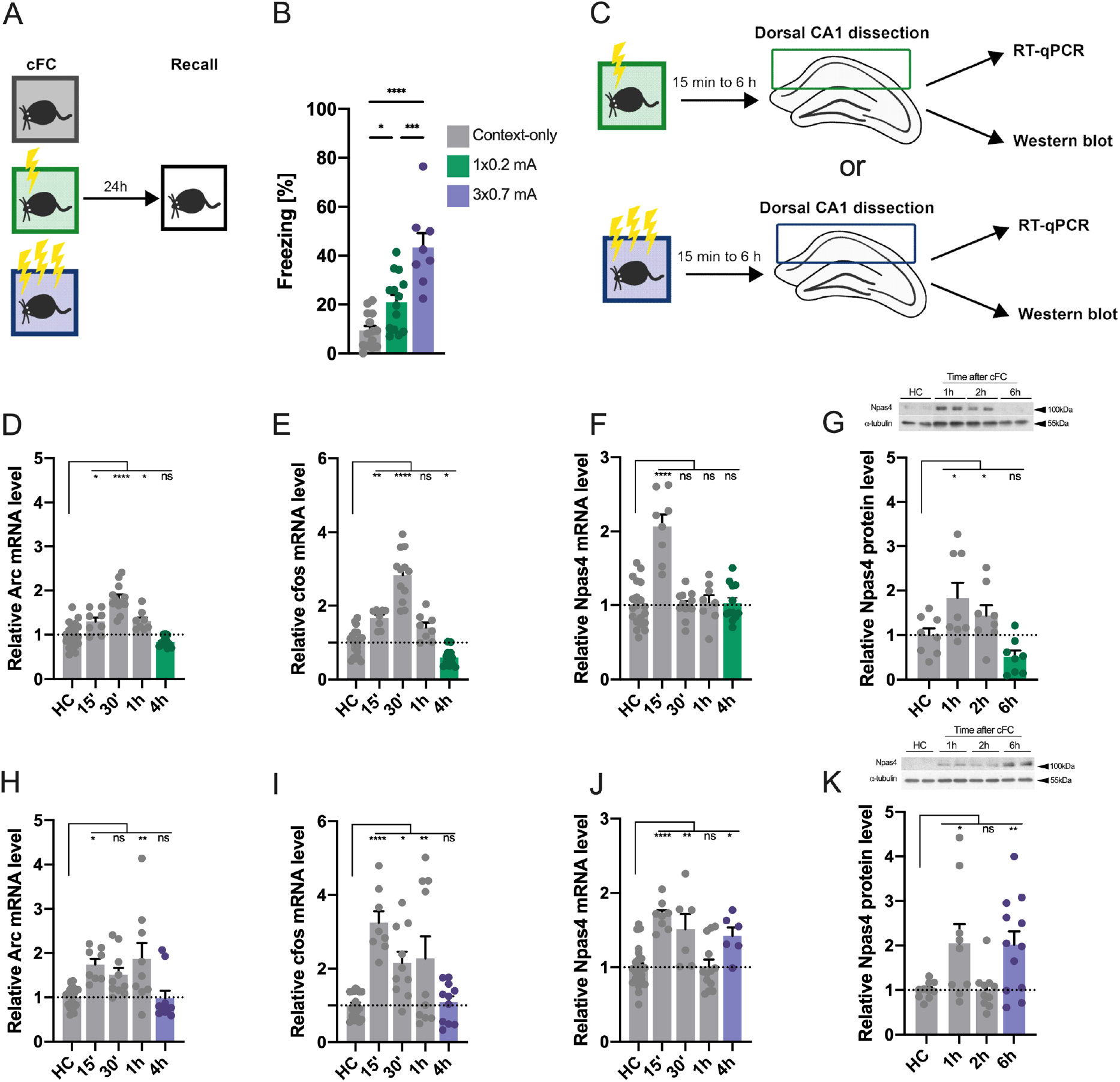
High salience contextual learning induces a biphasic expression of Npas4. (**A**) Experimental scheme. (**B**) Freezing behavior of mice exposed to a contextual fear conditioning (cFC) chamber without shock administration (context-only, N=14), or low (1x 0.2 mA, N=14) or high (3×0.7 mA, N=8) salience fear conditioning and tested 24h later for long-term memory. (**C**) Experimental scheme. Mice were trained in low or high salience fear conditioning and were sacrificed 15min, 30 min, 1h, 2h, 4h or 6h thereafter. Their hippocampi were dissected and mRNA or protein was isolated to perform RT-qPCR or western blot, respectively. Tissue from home cage (HC) control mice was collected at the same time of day to control for circadian alterations. mRNA levels of (**D**) Arc (N=8-20), (**E**) cfos (N=8-20) or (**F**) Npas4 (N=8-20) in mice that underwent low salience fear conditioning. (**G**) Protein levels of Npas4 (N=8) after low salience fear conditioning. mRNA levels of (**H**) Arc (N=8-20), (**I**) cfos (N=8-20) or (**J**) Npas4 (N=6-20) in mice that underwent high salience fear conditioning. (**K**) Protein levels of Npas4 (N=10-12) after high salience fear conditioning. Dots represent individual mice. Data represents mean ± standard error of the mean (SEM). One-way ANOVA; Dunnett’s or Šídák’s test; Ns, nonsignificant, *p<0.05, ** p<0.01, *** p<0.001, **** p<0.0001.

### NMDA receptor activity regulates late Npas4 expression and high salience contextual learning

The late Npas4 expression occurred several hours after the initial learning session, which raises at least two possibilities. First, the high salience stimulus might initiate a cascade of molecular events that *per se* would lead to a cell autonomous expression of Npas4, without the requirement for extracellular stimuli. An alternative hypothesis would be that several hours after learning, neurons receive new inputs that will lead to the late Npas4 expression. Previous studies showed that Npas4 expression is coupled to neural activity and dependent on NMDAR function and calcium influx.

Furthermore, IEG late expression events have been shown to be regulated by neuromodulatory transmitters, such as dopamine ^35, 36^. Therefore, we tested whether NMDAR or dopaminergic activity may trigger the second Npas4 expression wave in high salience fear conditioning. To this end, freely moving mice that underwent high salience fear conditioning were subsequently infused with the NMDA receptor antagonist - APV or the dopaminergic antagonist - SCH23390, 3.5h after training into the CA1 to evaluate the impact on the late Npas4 expression (Figure 2A). We found that blocking dopaminergic function had no effect on late protein levels of Npas4, Fos or Arc compared to saline infused animals (Figure 2B-D). In contrast, inhibiting NMDAR function decreased Npas4 protein levels at 6h (Figure 2B), whereas Fos and Arc levels were unaffected (Figure 2C-D). These findings are in line with previous studies that showed Npas4 expression to be highly dependent on neuronal activity and calcium influx, but not induced by signaling pathways that regulate expression of other key IEGs such as c-Fos and Arc, through growth factors or dopaminergic activation ^37, 38^. Taken together, this indicates that late Npas4 expression is dependent on NMDAR activity.

**Figure 2.**
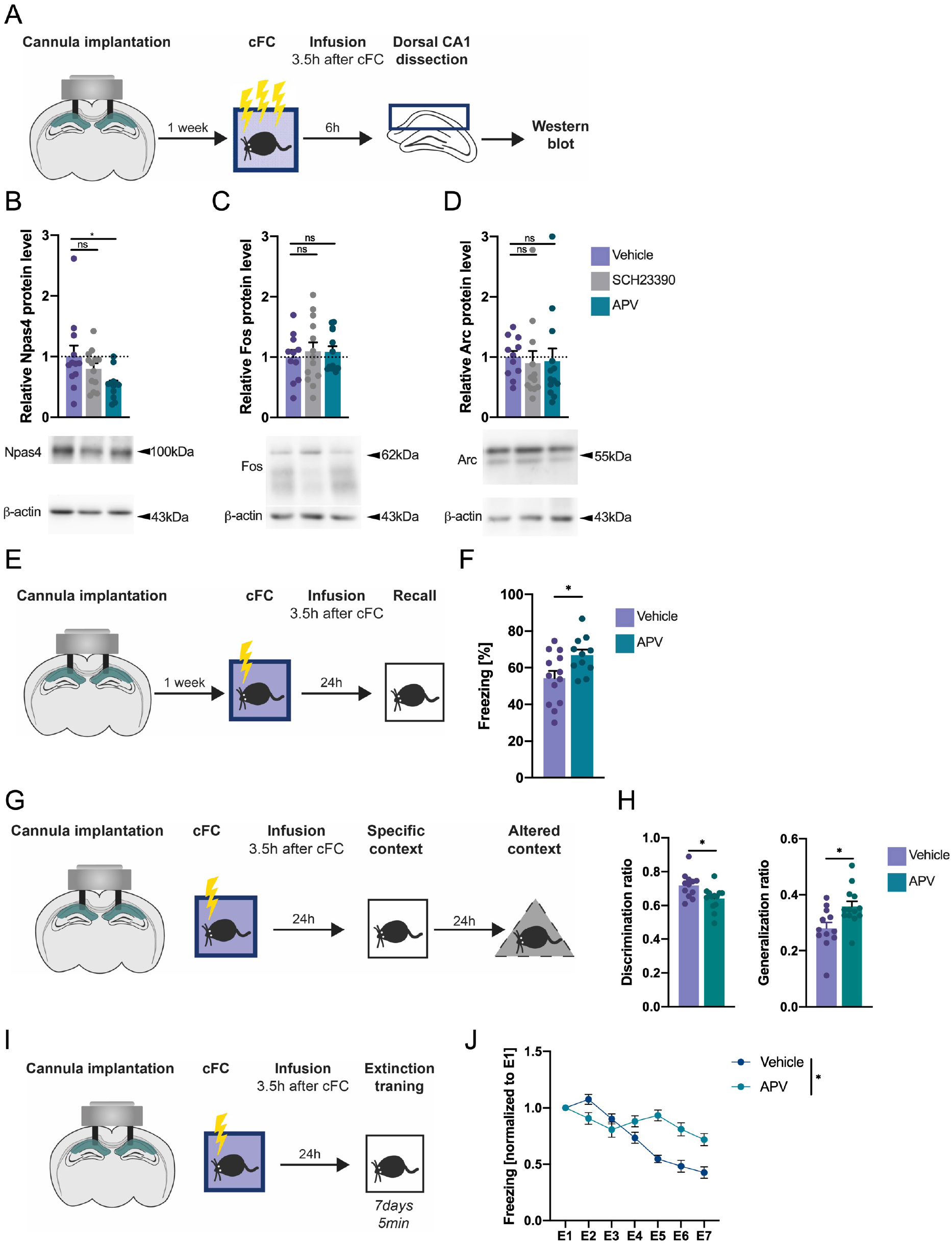
NMDA but not dopamine receptor activity regulates late Npas4 expression and constricts high salience contextual memory and maladaptive behavior. (**A**) Experimental scheme. 1 week after cannula implantation, mice underwent high salience contextual fear conditioning (cFC) training and 3.5h later mice were infused with vehicle or pharmacological inhibitors of dopamine D1 receptors (SCH23390) or NMDA receptors (APV). 2.5h later (i.e., 6h after training) hippocampi were collected and protein isolated for western blot analysis. (**B**) protein levels of Npas4 (N=11-12), (**C**) Fos (N=11-12) or (**D**) Arc (N=11-12). (**E-F**) Experimental scheme (**E**) and behavioral analysis (**F**) of LTM after NMDAR blockade. A cannula targeting the CA1 region of the hippocampus was implanted. 3.5h after high salience fear conditioning training, mice were infused with vehicle or APV and 24h after tested for long-term fear memory (N=11-13). **(G-H)** Experimental scheme **(G)** and behavioral analysis **(H)** of context discrimination and generalization. A cannula targeting the CA1 region of the hippocampus was implanted. 3.5h after high salience fear conditioning training, mice were infused with vehicle or APV and 24h after tested for long-term fear memory, 24h after this they were exposed to an alternate context and freezing levels were scored. A discrimination (specific context/[specific context + altered context]) and generalization (altered context/[specific context + altered context]) ratios were calculated (N=12-13). **(I-J)** Experimental scheme **(I)** and behavioral analysis **(J)** of extinction. A cannula targeting the CA1 region of the hippocampus was implanted. 3.5h after high salience fear conditioning training, mice were infused with vehicle or APV and during 7 consecutive days they were exposed to the same context for 5min to promote fear extinction (N=12). Dots represent individual mice. Data are shown as mean ± standard error of the mean (SEM). One-way ANOVA; Šídák’s test, two-way ANOVA, and unpaired Student’s T test; Ns, nonsignificant, *p<0.05.

Next, we asked whether NMDAR activation, which drives late Npas4 expression, regulates memory consolidation. To avoid possible confounds of overtraining, we established a second high salience fear conditioning protocol that also induces Npas4 biphasic expression (1×0.7 mA; Supplementary Figure 1). Mice trained in this protocol received APV or vehicle infusions into CA1 and were tested for memory performance 24h later (Figure 2E). Blocking NMDAR activity, and consequently late Npas4 expression, resulted in enhanced LTM compared to vehicle infused animals (Figure 2 F). Strong and maladaptive fear memories trigger disproportionate and generalized fear responses that are also resistant to extinction ^39, 40^. Thus, we investigated whether inhibition of the late NMDAR activity may prevent maladaptive responses namely the generalization of fear to a neutral context (Figure 2G,H) and the prevention of fear extinction (Figure 2I,J). We found that mice that received APV into the CA1 exhibited poor discrimination between the specific context (context in which the shock was delivered) and a neutral context compared to vehicle-treated mice (Figure 2H). Moreover, APV-treated mice showed impaired fear memory extinction (Figure 2J). These experiments show that late NMDAR activity suppresses the consolidation of highly salient fearful experiences and prevents maladaptive responses.

### Biphasic Npas4 expression suppresses memory consolidation

To directly assess the potential role of late Npas4 expression as an inducible memory suppressor gene, we took advantage of a Tet-On system to induce biphasic Npas4 expression. In order to model the induction of Npas4 expression at the time point that occurs physiologically in a high salience fear conditioning paradigm, we used recombinant adeno associated viruses (rAAV) harboring the Tet-On system *in vivo*. For tight control of tet-dependent transgene expression in neurons, one viral construct contained the human synapsin (hSyn) promoter expressing the reverse tetracycline-controlled transactivator (rtTA) together with the Tet repressor (TetR). The second construct contained tet-dependent HA-tagged GFP (GFP) or full-length HA-tagged Npas4 (Npas4) transgenes. Administration of doxycycline to hippocampal primary cultures verified tight control of transgene expression via the Tet-On system (Supplementary Figure 2A-C). To evaluate the expression kinetics of this system *in vivo*, we delivered the viral vectors into the CA1 region and monitored transgene expression (Supplementary Figure 2D). Exogenous Npas4 protein was detectable 6 hours after a single intraperitoneal doxycycline administration and peaked at 24h (Supplementary Figure S2D-G). Next, we confirmed the functional activity of exogenous Npas4 expression (Supplementary Figure 2H-I). Primary hippocampal cultures were co-transfected with these constructs (GFP or Npas4) and reporter plasmids that contain luciferase under the control of Npas4 responsive elements (NRE) or CREB responsive element (CRE) as a control and a minimal promoter (Supplementary Figure 2H). This assay reveled that exogenous Npas4 expression selectively increases NRE-associated luciferase activity (Supplementary Figure 2I). Altogether, this set of experiments demonstrated that our approach with the Tet-On system allowed time-dependent expression of functionally active Npas4 in hippocampal neurons.

To understand how artificial induction of late Npas4 expression would influence memory consolidation, we delivered rAAVs expressing GFP or Npas4 into the CA1 of mice and trained them in the low salience fear conditioning paradigm (Figure 3A). Given our preliminary observations showing that doxycycline treatment induces higher freezing levels compared to saline treated mice (data not shown), in this set of experiments both the control (GFP) and Npas4 groups received intraperitoneal doxycycline administration. In order to express Npas4 protein at the timepoint in which it occurs in high salience contextual learning (6 hours after training) (Supplementary Figure 2E), we administered doxycycline immediately after contextual fear conditioning (Figure 3A). This approach is limited by the lack of control of transgene expression levels. However, despite this limitation we achieved at 6 h after training, expression levels of exogenous Npas4 protein (Supplementary Figure 2E) that are within the same order of magnitude of the endogenous (Figure 1K) (7.8 fold ± 2.2 (mean ± SEM) versus 2.0 fold ± 0.3 (mean ± SEM) above control conditions for exogenous and endogenous, respectively). Mice that expressed late Npas4 within the window of memory consolidation displayed LTM impairments compared to the GFP control group (Figure 3B). This result suggests that biphasic Npas4 expression constricts memory consolidation.

**Figure 3.**
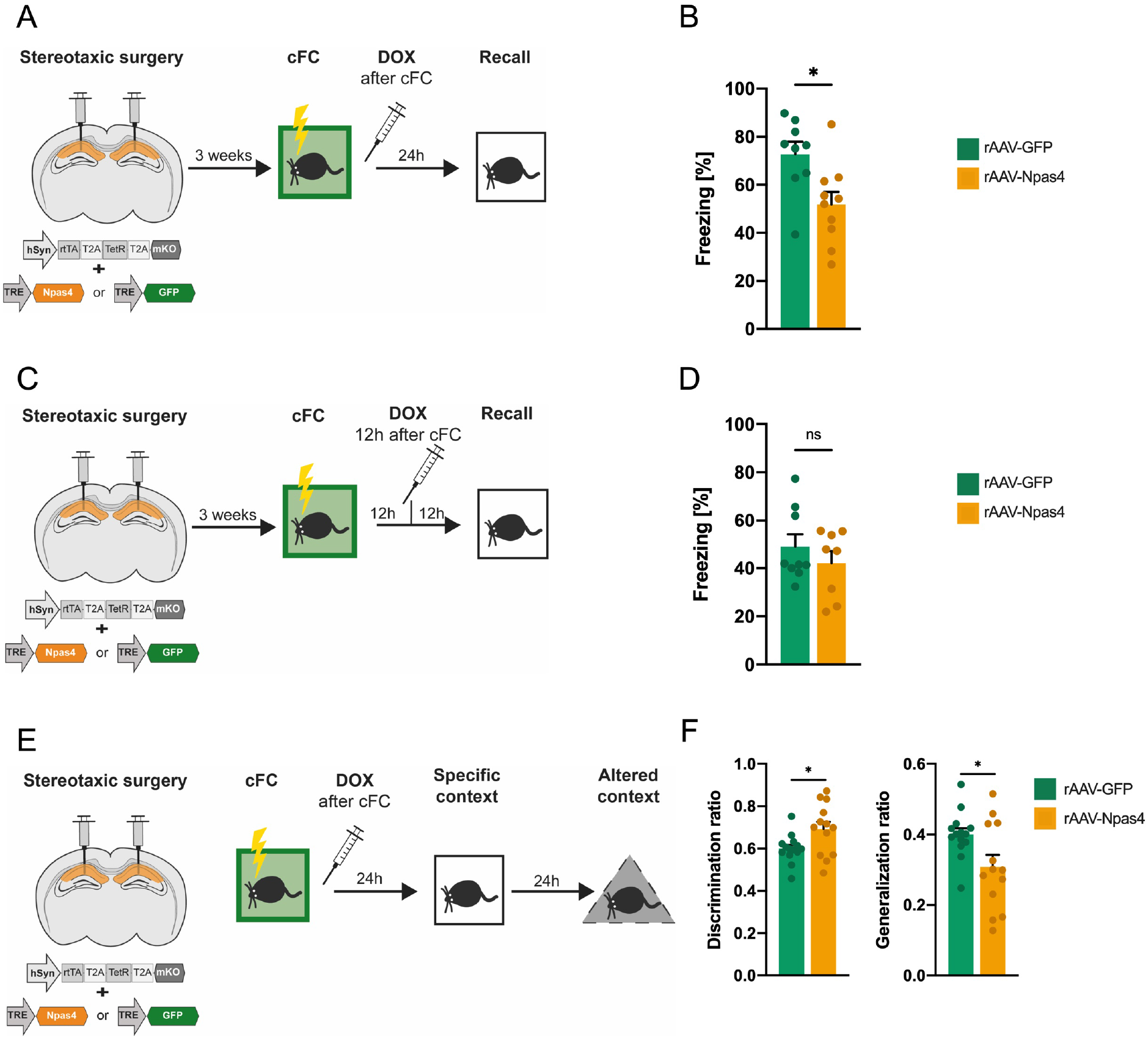
Late Npas4 expression suppresses memory consolidation and maladaptive behavior. (**A**) Experimental scheme. rAAVs expressing GFP or Npas4 were delivered to the CA1 region of the hippocampus. 3 weeks after surgeries mice underwent low salience contextual fear conditioning (cFC) and immediately received a single doxycycline (DOX) injection to activate the expression of exogenous proteins. (**B**) Behavioral analysis of LTM 24h after cFC training(N=9-10). (**C**) Experimental scheme. rAAVs expressing GFP or Npas4 were delivered to the CA1 region of the hippocampus.3 weeks after surgeries mice underwent low salience contextual fear conditioning. 12h after training mice received a single doxycycline injection to activate the expression of exogenous proteins. (**D**) Behavioral analysis of LTM 24 after training (N=8-9). (**E**) Experimental scheme. rAAVs expressing GFP or Npas4 were delivered to the CA1 region of the hippocampus. 3 weeks after surgeries mice underwent low salience contextual fear conditioning and immediately received a single doxycycline injection to activate the expression of exogenous proteins and 24h later were tested for long-term fear memory. 24h after this they were exposed to an alternate context and freezing levels were scored. **(F)** A discrimination (specific context/[specific context + altered context]) and generalization (altered context/[specific context + altered context]) ratios were calculated (N=13-14). Dots represent individual mice. Data are shown as mean ± standard error of the mean (SEM). Unpaired Student’s T test; Ns, nonsignificant, *p<0.05.

To evaluate whether increased Npas4 expression was specifically necessary at 6h after training for impairment, we temporally shifted exogenous Npas4 expression. Using a similar experimental design, we induced expression of Npas4 at a later stage (12h), which did not overlap with the 6h time point but is still within the memory consolidation phase ^11^(Figure 3C). We found that expressing Npas4 outside this specific time window had no impact on LTM formation, further indicating that the memory suppressor function of Npas4 is coupled with the 6h timepoint but does not appear to impact memory consolidation at a later time point (Figure 3D). These results demonstrate that the induction of Npas4 expression 6h after fear conditioning learning impairs memory consolidation, supporting its memory suppression function and highlighting a critical link between the temporal pattern of expression and this activity.

Lastly, to further strengthen a possible link between Npas4 late expression and the regulation of maladaptive responses, we investigated how artificial induction of late Npas4 expression influences the generalization of a fear response (Figure 3E). To this end, we delivered rAAVs expressing GFP or Npas4 into the CA1 of mice and trained them in the low salience fear conditioning paradigm. The mice freezing responses were later tested in a specific and a neutral context (Figure 3E). We found that mice that expressed late Npas4 displayed improved contextual discrimination compared to the GFP control group (Figure 3F). Taken together these findings further support the second Npas4 expression wave as an important molecular mechanism in the regulation of fear memory strength and maladaptive behavior.

### Biphasic Npas4 expression promotes inhibition from CCK^+^-expressing interneurons

Previously, it was shown that Npas4 expression regulates the inhibitory drive onto hippocampal pyramidal neurons ^41^. Specifically, Npas4 enhances the component of perisomatic inhibition provided by CCK^+^ IN ^20, 30^. Thus, we investigated whether the late Npas4 expression modulates CCK^+^IN-mediated inhibition by evaluating the connectivity between CCK^+^ interneurons and CA1 pyramidal cells in animals trained with the high or low salience fear conditioning protocol (Figure 4A). To detect CKK^+^ inhibitory synapses within the CA1 region we used immunofluorescence analysis of the presynaptic vesicular GABA transporter (VGAT), the inhibitory postsynaptic scaffolding protein gephyrin and the presynaptic cannabinoid receptor (CB1R) (Figure 4B)^20^. Upon quantification of the triple overlap, we found increased immunofluorescence signal, which was used as a proxy for CCK^+^ specific inhibitory synapses, in the superficial layer of the CA1 in tissue from mice that underwent the strong fear conditioning protocol (Figure 4C). Importantly, this regional specificity is in line with previous studies that reported selective Npas4-dependent regulation of inhibitory synapses located in the superficial but not deep CA1 layer ^20^. Next, we analyzed the inhibitory currents on acute hippocampal slices prepared from mice that underwent either the strong or weak fear conditioning protocol (Figure 4D,E). Twenty-four hours after training, CCK^+^IN-mediated inhibition was assessed using the unique property of these interneurons-sensitivity to endocannabinoids. Binding of endocannabinoids to the cannabinoid receptor type 1 (CB1R) expressed on presynaptic terminals of CCK^+^ neurons almost entirely blocks synaptic vesicle release ^42^. Synthesis of endocannabinoids can be triggered by depolarization of the postsynaptic cell. Hence, mIPSCs recorded at 0 mV should arise from the activity of all other interneurons except CCK^+^IN, while subsequent blockade of CB1R should reveal the actual contribution of CCK^+^IN to the net inhibition. We found that in slices from animals trained with the strong fear conditioning protocol, application of the CB1R blocker - AM-251 – resulted in a significant increase in frequency of mIPSCs (Figure 4E). In contrast, in mice exposed to the low salience protocol, AM-251 did not have an effect on mIPSCs (Figure 4D). Thus, strong but not weak fear conditioning results in increased inhibitory synaptic input to CA1 pyramidal neurons from CCK^+^ IN.

**Figure 4.**
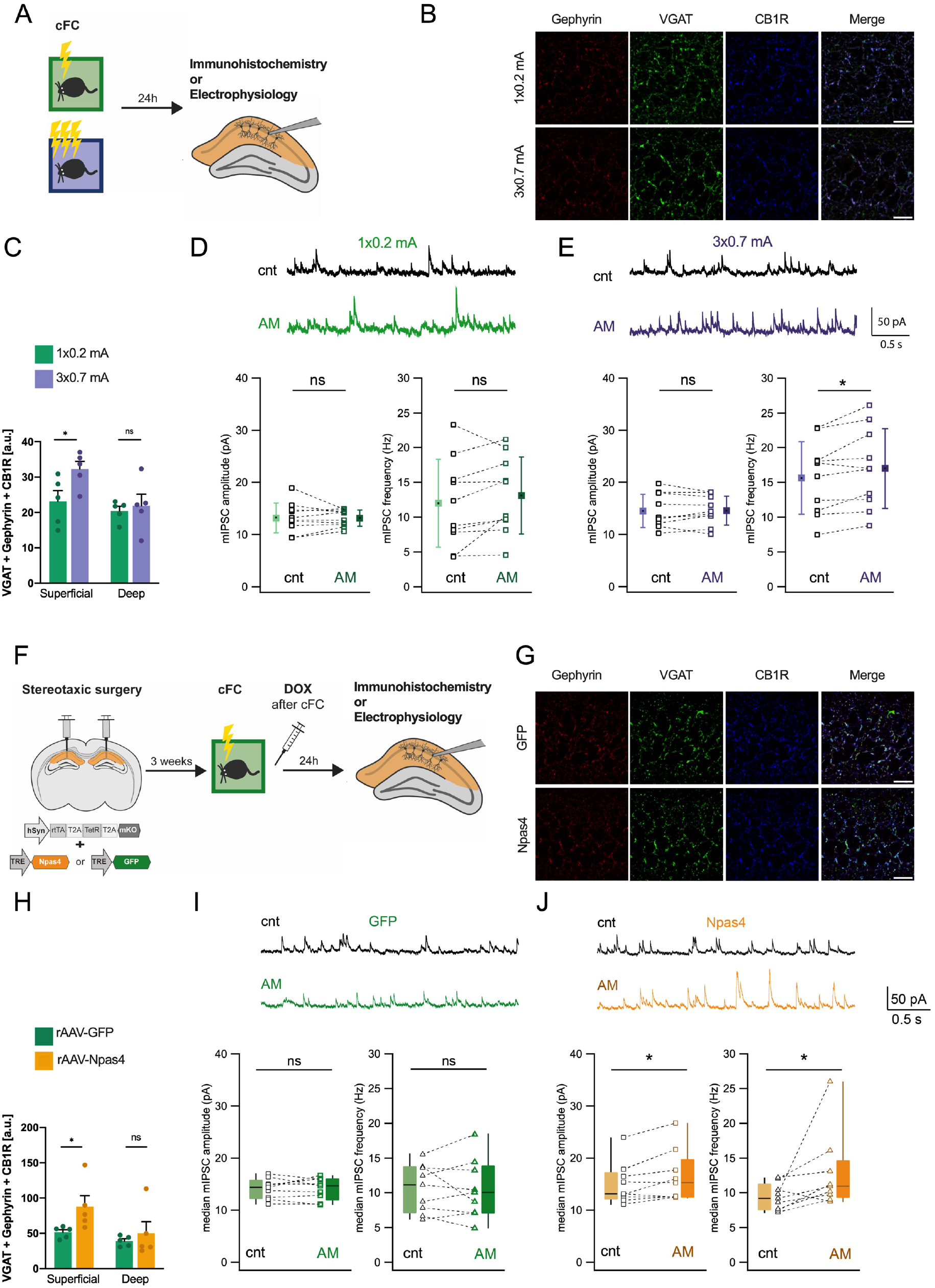
High salience fear conditioning and late Npas4 expression enhance inhibition from CCK expressing interneurons prior to memory recall. (**A**) Experimental scheme for data shown in B-E. Mice underwent low or high salience fear conditioning training and 24h later were sacrificed and their brains were processed for immunofluorescence staining or electrophysiology. (**B**) Representative confocal images of the CA1 region immunolabelled for gephyrin, VGAT and CB1R. Scale bar = 15 μm. (**C**) Quantification of colocalization in the superficial and deep regions of the pyramidal layer to identify synapses between cholecystokinin-expressing interneurons (CCK+ IN) and pyramidal neurons (n=5 mice). Dots represent average signal from sections for each individual mice. Data are shown as mean ± standard error of the mean (SEM). Unpaired Student’s T test, Ns:not significant, *p<0.05. **D-E**) Representative whole cell recordings showing mIPSCs in CA1 pyramidal cells before (cnt) and after CB1R blockade with AM-251 (AM) from animals trained with the low (**D**) or high (**E**) salience paradigm. Histograms show the median values for amplitude and frequency of mIPSCs in individual neurons (dots) before and after CB1R blockade. Average data show the mean ± SD. Significance of the effects of AM251 were assessed by T-test (Ns:not significant; *p<0.05; N=10 cells from 6 mice per group). (**F**) Experimental scheme for data shown in G-J. rAAVs expressing GFP or Npas4 were delivered to the CA1 region of the hippocampus. 3 weeks after surgeries mice underwent low salience contextual fear conditioning and immediately received a single doxycycline injection to activate the expression of exogenous proteins. 24h later were sacrificed and their brains were processed for immunofluorescence staining or electrophysiology. (**G**) Representative confocal images in the CA1 region for gephyrin, VGAT and CB1R. Scale bar=15 μm. **(H)** Quantification of colocalization in the superficial and deep regions of the pyramidal layer to identify synapses between CCK+ IN and pyramidal neurons (n= 5 mice). Data are shown as mean ± standard error of the mean (SEM). Unpaired Student’s T test, Ns:not significant, *p<0.05. Dots represent average signal from sections for each individual mice. **(I-J)** Representative whole cell recordings of mIPSCs in CA1 pyramidal cells expressing either GFP (N=9 cells from 4 mice) (**I**) or Npas4 (N=11cells from 4 mice) (**J**) before and after AM-251 application. Histograms show the median amplitude and frequency of mIPSCs in individual neurons (dots) before and after CB1R blockade. Since data on mIPSC frequency in Npas4 expressing mice did not have normal distribution, the pooled data presented as box plots show the median (line), 25th, 75th percentiles (box) and minimum and maximum (whiskers). Significance of the effects of AM251 were assessed by Wilcoxon Signed Rank Test; (Ns: nonsignificant, *p<0.05).

Next, to further support these findings, we assessed whether exogenous Npas4 expression modulates inhibitory synapses. We delivered rAAVs expressing GFP or Npas4 into the CA1 of mice and trained them in the low salience fear conditioning paradigm. Immediately after training, doxycycline was administered and 24 hours later slices were collected for either immunofluorescence analysis or electrophysiological recordings (Figure 4F). The analysis of the overlap of VGAT, gephyrin and CB1R immunostaining (Figure 4G) revealed that Npas4 overexpression led to a significant increase in immunofluorescence in superficial CA1 compared to control (Figure 4H). In order to determine whether this change in immunofluorescence of CCK^+^ inhibitory synapses translates into changes in functional connectivity, we measured mIPSCs (Figure 4I,J). We found that in CA1 pyramidal cells infected with rAAV-Npas4, application of the CB1R blocker - AM-251 - resulted in a significant increase in both amplitude and frequency of mIPSCs (Figure 4J). In slices obtained from control animals, AM-251 did not have an effect on mIPSCs (Figure 4I). Taken together, these observations support the notion of a selective enhancement of CCK^+^IN-mediated inhibition by late Npas4 expression.

## Discussion

In this study, we uncovered a novel mechanism regulated by the salience of a fearful experience that gates the strength of fear memory and associated maladaptive responses (Figure 5). We showed that, unlike low salience experiences, highly salient experiences induced two phases of Npas4 expression. Using pharmacological and genetic approaches we showed that the late Npas4 expression constraints the consolidation of fear memory and prevents the formation of maladaptive behavior, namely the contextual generalization of the fear response and resistance to suppression by extinction. Finally, we found that this effect is associated with increased CCK^+^ IN-dependent inhibitory input onto CA1 pyramidal neurons.

**Figure 5.**
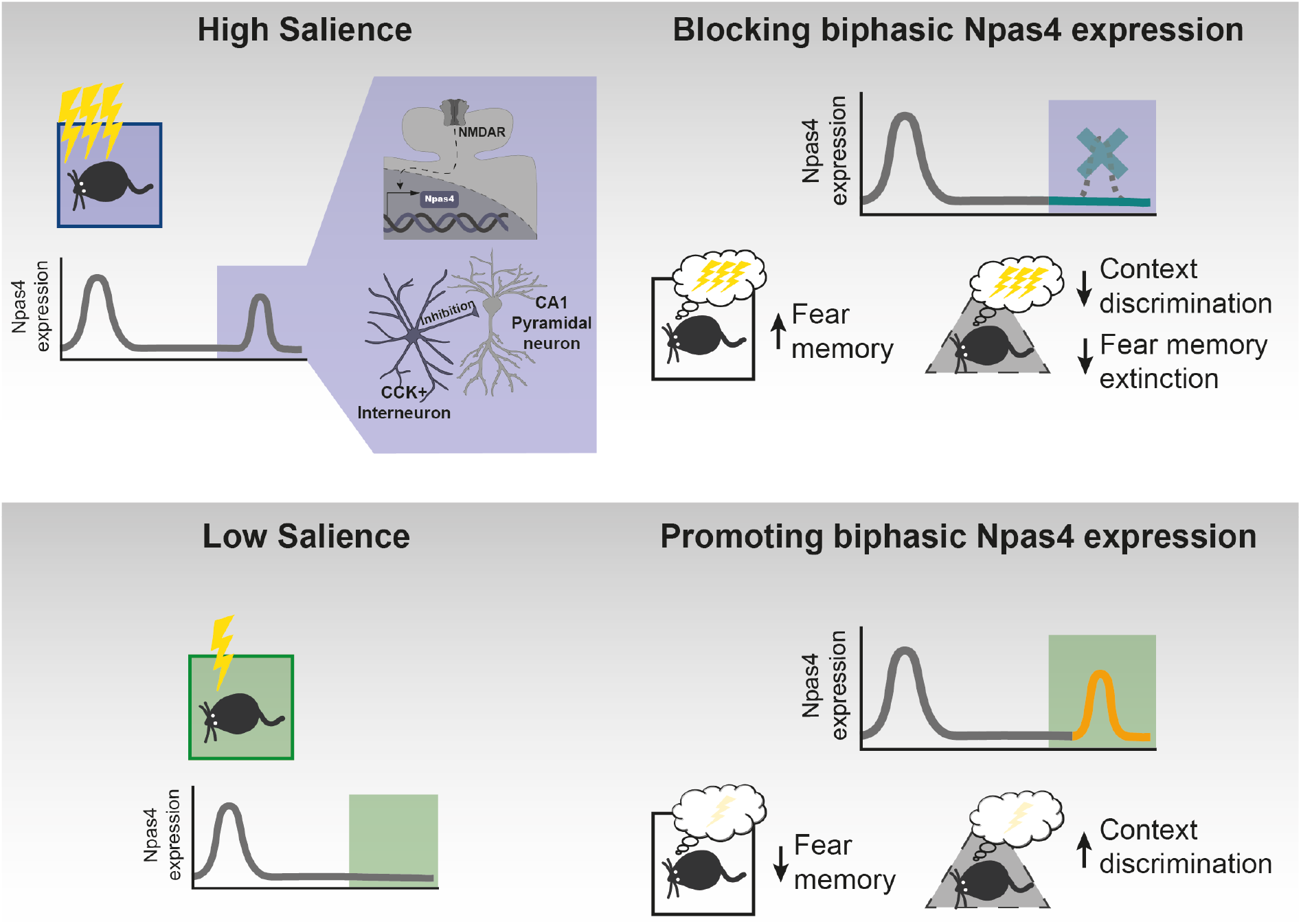
Graphical illustration of the main findings. Role of Npas4 biphasic expression for memory consolidation. Upper part: High salience learning (purple) induces Npas4 biphasic expression in the CA1 region of the hippocampus in a NMDAR-dependent manner and increases CCK^+^ IN inputs. Pharmacologically blocking NMDAR activity, and consequently late Npas4 expression, results in enhanced fear memory, reduced context discrimination and impairs fear memory extinction. Nonetheless, it should be noted that other mechanisms besides Npas4 expression occur due to NMDAR activation and may contribute to memory suppression. Lower part: Low salience learning (green) leads to one peak of Npas4 expression. Genetically promoting a biphasic expression of Npas4 results in reduced fear memory and increased context discrimination.

Learning experiences have been proposed to be encoded into long-term memory by genomic responses ^18, 34, 43^. Interestingly, the expression pattern of activity-regulated genes appears to define aspects of the learning experience. One study evaluated 13 different experiences with distinct attributes, such as salience and valence, and found that they are represented by unique transcriptional signatures ^15^. Analysis of the expression patterns of five IEGs across five brain regions was sufficient to predict with high efficiency the experience of individual mice. Besides experience attributes, the duration of neuronal activity has been shown to modulate the gene expression profile ^18^. Brief and sustained activity bursts induced an early genomic response, while prolonged neuronal activity generated multiple additional waves of gene expression. Furthermore, several studies have reported additional waves of gene expression associated with experience salience that take place hours after the initial experience. Particularly, highly salience conditioning was shown to induce multiple waves of IEG expression in mice, such as cfos, brain-derived neurotrophic factor (bdnf) and the early growth response 1 (egr1) gene ^11–14^. These studies demonstrated that the delayed expression wave of these IEGs promotes the persistence of memory. Here, we found that late IEG expression may also function to constrain memory consolidation. We discovered that the expression pattern of the IEG Npas4 is regulated by experience salience. Our results showed that highly salient experiences induced two phases of Npas4 expression and that the late expression phase suppresses memory consolidation and is regulated by NMDAR activity. A previous study evaluated the contribution of NMDAR activity for post-acquisition memory consolidation ^44^. Interestingly, the authors observed that rats treated with NMDAR antagonists 6h after learning displayed enhanced LTM in a spatial memory paradigm. These findings agree with our results that NMDAR blockade led to enhanced LTM and further suggest that the delayed wave of Npas4 expression may be a NMDAR-dependent mechanism that suppresses memory consolidation of highly salient experiences. Nonetheless it is probable that other mechanisms besides Npas4 expression occur due to NMDAR activation and contribute to memory suppression. Despite several attempts, the implementation of a method to achieve the time-specific abolishment of Npas4 second expression period was not successful. However, our gain-of-function approach provides compelling evidence that Npas4 expression from 6h after learning, and not at later time points, limits memory consolidation.

One of the leading hypotheses on the relevance of memory suppressor genes is to possibly allow behavioral flexibility ^6^. For example, without constrains on consolidation, highly salient experiences might generate strong memories that are not adaptive or advantageous in other scenarios with partially overlapping contexts and cues. Taken together, this and previous studies suggest that stimulus salience is encoded by an IEG expression pattern which regulates not only whether an experience will be converted into LTM, but also modulates the strength of the memory. Furthermore, our study showed that highly salient experiences induce two phases of Npas4 expression with potentially opposing effects on memory consolidation. It is well established that Npas4 is required for memory consolidation as demonstrated by studies that abolished its expression prior to acquisition and throughout consolidation ^19, 37, 45, 46^. In our study, we found that Npas4 expression 6h post learning constrained memory consolidation, acting as an inducible memory suppressor gene, contrary to the early Npas4 induction wave. In this scenario, we propose that the first wave is likely required for the formation of memory, while the second wave fine tunes its strength, potentially preventing the formation of strong maladaptive memories. In support of this, we showed that NMDAR blockade prior to the second expression wave of Npas4 resulted in a stronger fear memory characterized by generalized fear responses and resistance to suppression by extinction. Conversely, the artificial induction of the Npas4 late expression resulted in increased contextual specificity of the behavioral response. In line with our findings, a recent study has shown that Npas4 regulates memory discrimination^30^.

However, how the Npas4 first and second expression phases lead to distinct effects on memory consolidation remains to be understood. Recent findings reported a role of Npas4 expression as a stimulus decoder in the brain ^47^. Particularly, inducing action potential or excitatory postsynaptic potentials in the CA1 led to the formation of stimulus-specific Npas4 heterodimers that bind to different genomic loci. This mechanism illustrates how different neuronal stimulations can be encoded by the same IEG. It is tempting to speculate that Npas4 expressed immediately or 4h after learning might interact with differentially available partners. In this scenario, Npas4 heterodimers can induce a salience-specific gene response that might facilitate or restrict memory consolidation ^48^. Nonetheless additional open questions remain unanswered: what is the cellular process that drives NMDAR-dependent Npas4 induction and how does it selectively induce Npas4 expression and not other IEGs?

And which transcriptional program does Npas4 expression induce at this timepoint that constrains memory formation?

At a cellular and circuit level, Npas4 has an established role in the modulation of activity-dependent synaptic connections. It has been suggested to take part in negative feedback mechanisms and to be involved in synaptic homeostasis to maintain neural circuit balance ^45^. Consistent with this function, it has been shown that the level of Npas4 expressed in an excitatory neuron dictates the amount of inhibitory drive it receives ^49^. Overall, these studies indicate that Npas4 regulates inhibitory synapses to fine-tune neuronal circuits. More recently, it has been shown that the expression of Npas4 in CA1 and dentate gyrus principal neurons selectively increased inhibitory input made by CCK^+^ IN onto the excitatory neurons ^20, 30^. The present study provides further support to this function. Specifically, using both immunostaining and electrophysiological analysis, we found that the formation of strong fear memory is associated with increased inhibitory input to neurons of the CA1 superficial layer from CCK^+^ IN. This is in line with a previous study that showed that optically evoked CCK^+^ IN inhibition onto Npas4 expressing neurons of the dentate gyrus, assessed 24 hours after learning, was stronger in mice that underwent a stronger contextual fear conditioning paradigm compared to weaker conditioning ^30^. Furthermore, we found that genetically inducing late Npas4 expression increases CCK^+^ IN-dependent input onto CA1 superficial layer. Notably the sublayer specificity is in line with the predominant presence of CCK^+^ terminals in the superficial compared to the deep CA1 pyramidal layer^50^. Taken together, our findings suggest that the second Npas4 expression period may be a contributor to the observed increase in CCK^+^ IN-dependent inhibitory activity. Although these findings propose an intriguing novel mechanism by which the extent of consolidation of highly salient experiences into LTM may be modulated, a causal link between Npas4-dependent modulation of CCK^+^ IN inhibitory output and memory strength remains to be established. This opens the exciting opportunity to identify the so far unexplored function of CCK^+^ IN in modulating memory formation and its strength in future studies.

It is tempting to speculate that the delayed Npas4 expression might play a role in redistributing local inhibitory input promoting synaptic homeostasis during memory consolidation leading to a weakened memory trace. It has been proposed that the consolidation of salient experiences involves the renormalization of net synaptic strength ^51^, a process that goes in line with the described functions of Npas4 ^45^. In light of this theory, learning flexibility and new learning are compromised in the absence of cellular homeostasis restoration during the consolidation of memory, which in the particular case of emotionally charged experiences, could give rise to maladaptive and persistent fear memories that underlie psychiatry conditions i.e., post-traumatic stress disorder (PTSD). Overall, our study uncovered a biological mechanism that modulates the strength of memories of highly salient experiences, which may also play a role in regulating resilience to adverse life events.

## Acknowledgments

We thank Kübra Gülmez Karaca for critical comments on the manuscript, Benjamin Zeuch, Stefanos Loizou, Franziska Mudlaff, Carmen Leibold for technical support and Iris Bünzli-Ehret for the preparation of primary hippocampal cultures. This work was supported by the Deutsche Forschungsgemeinschaft (DFG) [grant numbers OL 437/1, OL 437/2, OL 437/3 and OL 437/4 to AMMO and BE 7081/2-1 to CPB], the Chica and Heinz Schaller foundation [fellowship and research award to AMMO], the Heidelberg Graduate Academy [Landesgraduiertenförderung (LGF) completion grant to DVCB]. AR and RS are supported by RSF 22-15-00293 and by “Center of Photonics” [funded by the Ministry of Science and Higher Education of RF (contract no. 075-15-2022-293)], respectively.

## Author Contributions

DVCB, JK, AMMO designed research; DVCB, JK, AR, CPB performed experiments; DVCB, JK, RS, MB, AR analyzed data; SC contributed new reagents; DVCB, AMMO wrote the paper. All authors edited the manuscript.

---

**Supplementary Figure 1.**
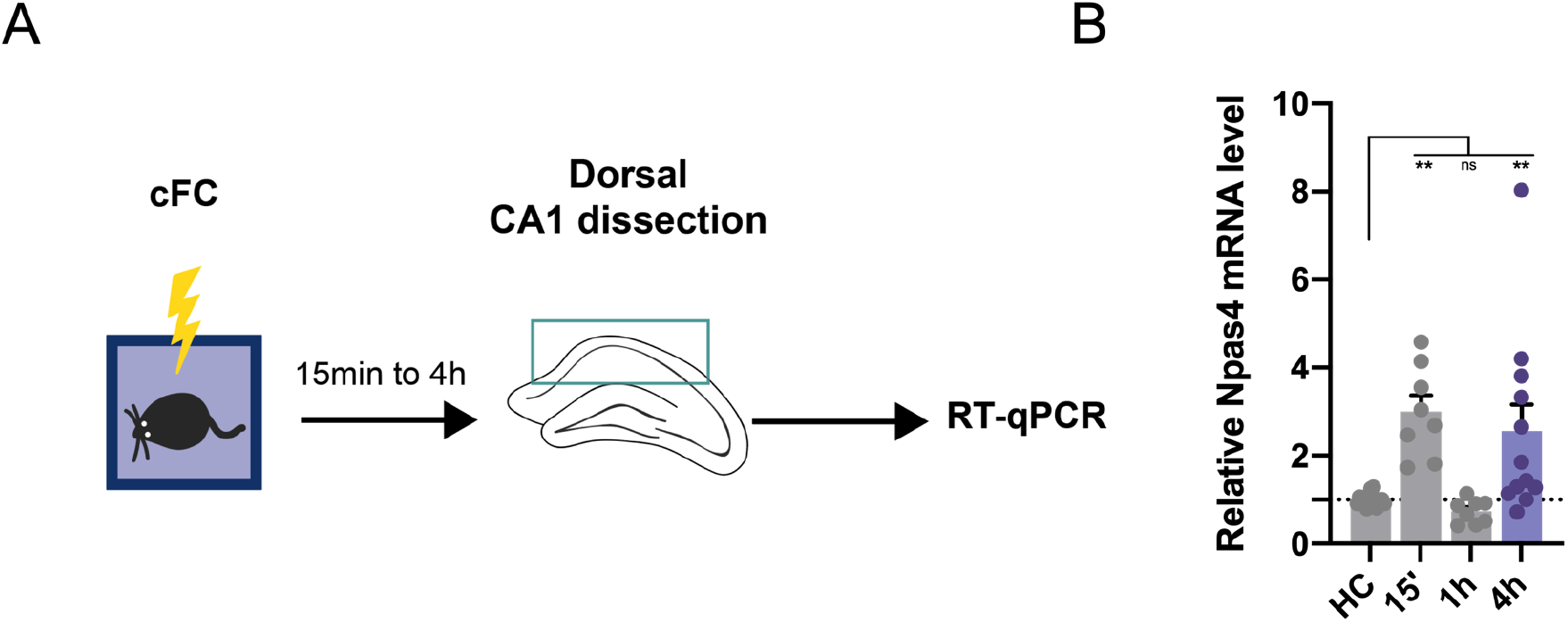
Validation of a second high salience contextual paradigm that induces Npas4 biphasic expression. **(A)** Experimental scheme. **(B)** qRT-PCR analysis of CA1 tissue obtained from mice that were left in their home cage (HC) or exposed to high (1×0.7 mA, N=8) salience fear conditioning training and sacrificed 15min, 1h or 4h after (N=8-15). Dots represent individual mice. Data are shown as mean ± standard error of the mean (SEM). One-way ANOVA; Dunnett’s; Ns, nonsignificant, ** p<0.01.

**Supplementary Figure 2.**
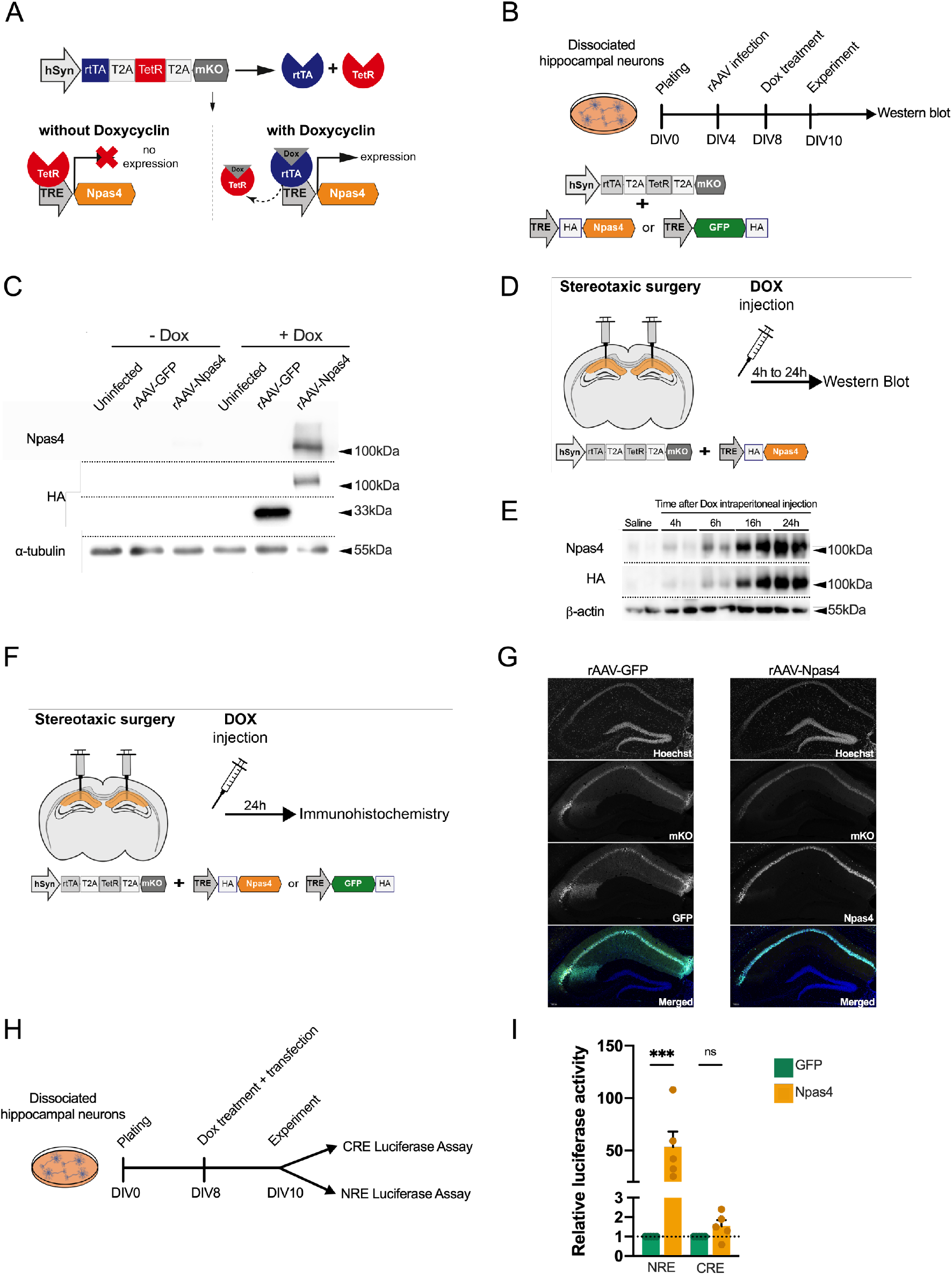
Expression and functional characterization of the dual-component TetON-based system. (**A**) Construct scheme. In the absence of doxycycline, the Tet repressor (TetR) is bound to the Tet response element (TRE) promoting active repression of transgene expression (left). In the presence of doxycycline, the TetR loses affinity thus allowing the reverse tetracycline-controlled transactivator (rtTA) to bind to the TRE and initiate transgene expression. (**B**) Experimental scheme. Primary hippocampal cultures were infected with rAAVs encoding the TetON-based system to express HA-tagged Npas4 or GFP. Doxycycline (Dox) was introduced at DIV8 to induce expression which was evaluated by **(C)** western blot using an antibody against Npas4 or the HA tag of exogenously expressed Npas4 or GFP. **(D)** Experimental scheme. rAAVs were delivered into the CA1 of mice and 3 weeks later mice received intraperitoneal injections of saline or Dox and sacrificed 4h, 6h, 16h or 24h later. **(E)** Expression of exogenous Npas4 was evaluated by western blot. **(F)** Experimental scheme. rAAVs were delivered into the CA1 of mice and 3 weeks later mice received intraperitoneal injections of Dox and sacrificed 24h later. **(G)** Expression of GFP or Npas4 was evaluated by immunohistochemistry. Scale bar= 100μm. **(H)** Experimental scheme. Primary hippocampal cultures were transfected with the dual-component TetON-based system and reporter plasmids that expressed luciferase under the control of a minimal promoter and Npas4 responsive elements (NRE) or CREB responsive element (CRE) and were treated with Dox (N=5 independent cell preparations). **(I**) Luciferase assay. Data are shown as mean ± standard error of the mean (SEM). Dots represent individual cell culture preparations. One-way ANOVA; Dunnett’s; Ns, nonsignificant, *** p<0.001.

